# Characterization of Mechanics Driven Heterogeneity in Mesenchymal Stromal Cells

**DOI:** 10.1101/2022.07.25.501486

**Authors:** Samantha Kaonis, Zack Aboellail, Soham Ghosh

## Abstract

Mesenchymal stromal or stem cells (MSC) are one of the most promising candidates for a myriad of cell therapy applications because of their multipotency, trophic properties and immunomodulatory properties. Despite showing promises in numerous preclinical and clinical studies, MSC based therapy is not yet a reality for regenerative medicine due their suboptimal outcome at the clinical endpoint. Suboptimal function of MSC is often attributed to the monolayer expansion process on plastic which is a necessary condition to reach the therapeutically relevant number, and also to their response to a fibrotic environment post transplantation. In both scenarios of plastic culture and fibrotic conditions, the mechanical environment experienced by the MSC is completely different from the natural mechanical niche of the MSC. Accordingly, the role of mechanical environment has been shown to be a critical determinant of MSC gene expression and function. In this study we report that human bone marrow derived primary MSC population becomes phenotypically heterogenous when they experience an abnormal mechanical environment, compared to their native environment. Using a newly developed technique to quantify the heterogeneity, we provide the evidence of phenotypical heterogeneity of MSC through high resolution imaging and image analysis. Additionally, we provide mechanistic insight of the origin of such substrate mechanics driven heterogeneity, which is further determined by the cell-cell mechanical communication through the substrate. The outcome of this study might provide mechanism driven design principles to the molecular, cellular and tissue engineering researchers for rational design of MSC culture condition and biomaterials, thus improving their functional outcome.

## INTRODUCTION

Multipotent mesenchymal stromal/stem cells (MSC) from different sources such as bone marrow and adipose tissues are promising candidates for tissue engineering and regenerative medicine applications [1]. MSCs are the candidate in more than a thousand FDA-approved clinical trials due to their multipotent differentiation ability, immunomodulatory properties, and trophic properties. Despite such optimism with MSC, the scalable application of MSC at the clinical endpoint is limited [2]. Autologous MSC therapies require a bone marrow aspirate from the patient. However, of the bone marrow mononuclear cells harvested, only 0.001% to 0.01% of the cell population comprises true MSCs [3]. To enrich the MSC content of the bone marrow aspirate, the cells are processed through density gradient centrifugation followed by direct plating [4]. Cells are deemed as MSC when they form the fibroblast colony-forming units (CFU-F) after plastic adherence [5]. Under these practices, only around 700 CFU-F/ml bone marrow aspirate can be collected [6], which is too small in comparison to the number of cells needed for therapeutic purposes, so expansion to millions of cells is required. For reference, between 8 and 20 million cells are required in the treatment of chondral defects using matrix-induced autologous implantation with MSCs [7,8], and for MSC therapies a dose of roughly about 100 million cells/70 kg are required to treat a patient if injected directly in the bloodstream [9].

The release criteria for MSCs is variable in clinical trial reports and is not standardized or required by the regulatory agencies such as FDA. It was reported that only 53.6% of clinical trials examined for the specific cell markers in the cell population, with assessment of functionality being very limited, sometimes falling short of the minimum suggestions by the International Society of Cell Therapy to ensure a population of MSCs [10,11]. Due to these inconsistencies, the cell population is not homogeneous at the outset of *in vitro* expansion, where other factors, such as seeding density, start to have an impact on the cells’ functional potential [12]. Subsequently, the *in vitro* expansion process to obtain such large numbers of cells becomes compromised by gradual differentiation of MSC into an increasingly larger population of osteoprogenitor like cells (non-MSC) that do not retain the MSC specific attributes, thus making the population heterogeneous. Even a small subpopulation of these “non-MSCs”, straying from the spindle-shaped population that generally represents multipotent cells [12], affect the population level functionality, limiting the therapeutic benefits at the clinical endpoint. Even when a cell population is established from a single MSC clone isolated in the culture, these cultures still become heterogeneous [13]. This reduced potential and increased heterogeneity can be attributed to the *in vitro* expansion on plastic as well as their response to the post-transplantation fibrotic environment. In the context of MSC culture, expansion and delivery in tissues, a key notable factor is that the mechanical environment on plastic or inside the fibrotic tissues are quite different from the natural soft niche experienced by MSC *in vivo* [14].

The substrate or the extracellular matrix mechanics of the cell affect the structure and mechanics of the cytoskeleton and the nucleus [15], which can have far-reaching impacts through gene expression and cell function, as shown in many cell types including MSC. Cells exert contractile forces on their culture substrate and the substrate elasticity can alter the migration behavior of endothelial cells, promoting the formation of cell-cell interactions [16] through integrins that transmit these mechanical signals via actin stress fibers and associated pathways [17]. Mechanical signal from the substrate elasticity can be relayed through the LINC (Link of Nucleoskeleton and Cytoskeleton) complex to the nucleus to trigger differential gene regulation pathways through epigenetic regulations [18] or through the engagement of transcriptional coactivators such as YAP (Yes-associated protein) and Transcriptional Co-Activator TAZ [17,19].

YAP/TAZ regulate key biological processes such as cell proliferation and migration, differentiation, and cell morphology [20] requiring Rho GTPase activity and tension exerted by the actomyosin contraction [21]. They shuttle between the cytosol and nucleus to regulate the target gene expression, being key mediators of cellular response to mechanical stimuli such as cell density, cell attachment area, shear forces, and substrate stiffness [22–24]. During the specification of MSC fate, YAP/TAZ can act as a molecular rheostat that modulates MSC differentiation [25]. On stiff substrates where the cells experience higher intracellular strain, YAP localizes to the nucleus and interacts with the transcription factor Runt-related transcription factor 2 (Runx2). Runx2 is the master gene of osteogenic differentiation [26], instructing the cells follow an osteogenic lineage which may potentially cause a heterogeneous shape and size distribution due to the deviation of the population away from the MSC phenotype. Conversely, on soft substrates there is lower intracellular strain and cytoskeletal F-actin formation, and accordingly MSCs favor MSC homeostasis, accompanied by down-regulation of Runx2 [27] and a more uniform cell shape and size [15]. This ability of MSCs to perceive stiffness and subsequently alter their phenotype can be influenced by cell seeding density as well. Contracting cells are known to transmit stress to neighboring cells over distances of tens of micrometers away, with the substrate stiffness being a key determinant in the ability of cells to mechanically communicate [28]. Cell seeding density can have a key role in the ability to transmit stresses through the substrate and cause changes in cell morphology, as shown in the migration of endothelial cells [16]. Cells that are seeded at a low density are known to show impaired proliferation or accelerated senescence in MSC. Therefore, understanding the mechanical interaction of cell clusters through substrate could be relevant to the MSC mechanosensitive response, population heterogeneity and function.

As cells replicate, differentiate, or senesce; measurable changes occur in the cell phenotype such as their size or the mechanical stiffness and these characteristics may serve as indicators of cell fate. Lee et al. (2014) examined multivariate biophysical markers and their potential to predict MSC multipotency and determined that a combination of small nucleus diameter, low cell stiffness, low F-actin content and high nuclear fluctuations suggestive of an open chromatin architecture, are indicative of undifferentiated subpopulations in culture-expanded MSCs [29]. During longer *in vitro* expansion, MSC nuclei became larger and the cell shape becomes wider and flatter with increasing morphological alterations [30,31]. These phenotype changes are indicative of an osteoprogenitor population, rather than a population of MSCs. Understanding the mechanisms of how the non-MSC subpopulation originates throughout *in vitro* culture can provide us with the ‘pre-treatment’ strategies to intervene the mechanisms to obtain a high-quality, homogeneous MSC population suitable for MSC-based tissue engineering/ regenerative medicine applications.

In this study we developed a technique to quantify the MSC population heterogeneity for investigating the mechanism of how the mechanical stiffness of the environment and the cell-cell mechanical communication through the growth substrate leads to a more heterogeneous population from undifferentiated MSCs. We hypothesize that neighboring cells communicate through compliant substrate, promoting a homogeneous population of cells with the MSC phenotype. To test this hypothesis, using human bone marrow derived primary MSC, we examined how substrate mechanics in combination with seeding density determines the MSC cell and nuclear phenotype over multiple passages, through high resolution imaging of the nucleus and the cytoskeleton. The functional meaning of population level MSC heterogeneity was assessed through the imaging of markers specific to MSC and Runx2. Knowledge of the mechanisms to maintain a homogeneous MSC population in two-dimensional culture can provide future intervention strategies to gain more control over MSC during their longer-term expansion in the clinical setting without requiring a specialized bioreactor system.

## MATERIALS AND METHODS

### MSC culture

Passage 2 primary human bone marrow derived MSC (PT-2501) were purchased from Lonza for all the experiments. These cells are mixed population of cells derived from the bone marrow of healthy adult individuals. Cells were maintained in bulletkit (Lonza) growth medium kit consisting the MSC Basal medium (PT-3238) and the supplemental kit (PT-4105) as stated in the manufacturer’s protocol. The culture condition was 5% CO2, 37°C temperature and 90% relative humidity. For all the experiments, cells were seeded in a 75 cm^2^ T-flask and passaged at 90% confluency at 5000 cells/ cm^2^, unless stated otherwise where in some cases cells were seeded at a higher or lower density. Trypsin-EDTA (0.05%) was used for subculturing. Medium was replenished every 3 days. Subsequently, passage 3 and 4 cells were used for all the experiments except for the cell treatment with the cytoskeleton modifying drug group where passage 5 cells were used. For all experiments, cells were directly seeded on the substrate after thawing the cells previously stored in liquid nitrogen tank.

### MSC culture on substrate with varying mechanical stiffness and cell density

Three substrate stiffness groups were prepared for cell culture. Soft: Sylgard 527 PDMS (*E*∼5 kPa), medium stiffness: Sylgard 184 PDMS (*E*∼1 MPa) and stiff: polymer resembling cell culture plastic (*E*∼1 GPa). Those mechanical stiffness conditions represent the physiological bone marrow stiffness, pathological fibrotic condition and the standard T flask cell culture condition respectively. Cells were cultured on 8 well chamber slides (ibidi, 80821) with thin bottom for high resolution imaging using a confocal microscope. For the stiff group, untreated polymer bottom slides were used followed by bovine Type 1 collagen coating (50 μg/ml, A10644-01, Gibco). For the soft and medium stiffness groups, the bottom polymer coverslip was removed, and custom-made chamber slides were refabricated with thin PDMS substrate as described in the next section. For the baseline cases, 5000 cells/cm^2^ cell density was used. To understand the effect of cell density on the MSC heterogeneity 100, 1000 and 10000 cells/ cm^2^ were used in addition to the 5000 cells/ cm^2^ group.

### Preparation of thin PDMS substrate

Polydimethylsiloxane (PDMS) substrates were prepared using Sylgard 527 and Sylgard 184 (Dow Corning). To fabricate the thin soft PDMS substrate (∼ 5 kPa), Sylgard 527 was mixed at a 1:1 ratio of each solution provided by the manufacturer. The mixture was then placed in a water bath at 37°C for 20-30 minutes to begin the curing process. Sylgard 184 thin PDMS sample was prepared by mixing ten parts of base with one part of curing agent and vigorous mixing. A small amount of each mixture (∼70 μl) was distributed across a glass coverslip (24×60 mm). The glass coverslips were then placed in a vacuum chamber for 20 minutes to remove any bubbles. For attachment of the coverslip to the bottomless chamber slide (ibidi, 80821), the coverslip was partially cured for 2 hours at 75°C and Sylgard 184 (10:1 mixture) was used to mount the coverslip to the chamber slide. To completely cure, it was placed in a 75°C oven overnight. Cured PDMS were then plasma treated (BD-20AC, Electro-Technic) to facilitate the collagen coating. Then they were sterilized in the cell culture cabinet with a 70% ethanol soak under UV light for 20 minutes, followed by incubation in a type I collagen coating (50 μg/ml, A10644-01, Gibco) for 1 hour to improve cell attachment and proliferation. The stiff group with polymer bottom was also treated with the plasma and the same type I collagen solution to maintain the same biochemical treatment between the groups. Cells were seeded on these prepared substrates and cultured with the previously stated culture condition.

### Cell Staining for immunofluorescence, actin and DNA

For all imaging tasks cells were fixed with 4% PFA (in 1X PBS: Phosphate Buffer Solution) for 10 minutes at room temperature. Cells were permeabilized with 0.1% Triton X-100 in 1X PBS, washed with 1X PBS, blocked for the non-specific binding sites with bovine serum albumin and normal goat serum. Primary antibodies for CD73 (41-0200) and Runx2 (PA5-82787) were then diluted at 1:100 in antibody dilution buffer and incubated overnight at 4°C. Next, secondary antibodies (A32731, A32727) were applied at 1:200 dilution at room temperature for 2 hours. All antibodies were purchased from ThermoFisher Scientific. Then, cells were stained for the DNA using DAPI (ThermoFisher Scientific) and for the F-actin, using phalloidin conjugated with fluorescent protein (ThermoFisher Scientific). Cells were imaged across their midsection using a confocal microscope (Zeiss LSM 980). Cells were imaged either at high resolution with 20X air objective or at low resolution with 10X air objective for all subsequent analysis. For all imaging, the imaging setting (laser power, gain etc.) was maintained constant between the groups. For the low-resolution imaging, the tile mode of imaging was used to image the complete surface area so that analysis could be performed to derive population level parameters in order to obtain a robust quantification of heterogeneity.

### Quantification of cell and nuclear morphology and stress fiber formation

#### Cell area and actin stress fiber quantification

To assess the amount of stress fiber in the cytoskeleton, the high resolution GFP-phalloidin stained images were used. Stress fiber intensity quantification was performed using a custom MATLAB code (Figure S1). Briefly, the code finds local maximum and minimum of the actin intensity and calculate that ratio as a measure of stress fiber formation. The same code was used to calculate the cell area. The cytoskeleton phenotype was not analyzed as a function of cell density effect due to technical limitations of identifying individual cells at the high seeding density.

#### Nuclear area and other geometrical parameters calculation

Nuclear morphology quantification was performed on the tiled images of the entire growth surface for a robust image analysis. Images were binarized and watershed separation was applied to separate cells that were touching or very close using ImageJ. These images were then imported into MATLAB. Subsequently, disk structuring elements were created through the strel() function, to create masks over each identified nucleus, which were then smoothed and filled to create a solid element. The size filtering for the nucleus images was critical to avoid any small background particles that may remain or any nuclei that were not separated by the watershed function. The final allowable elements were labeled for later reference and then measured with regionprops() function to calculate the nucleus area and circularity.

### Quantification of chromatin architecture

To calculate chromatin segregation from the DAPI stained nucleus images, a technique was adapted from a previous approach described by Irianto et al. [32]. The high-resolution images of nuclei were cropped and apportioned into images of individual nuclei. To calculate the chromatin segregation index based on the DAPI stain, gradient-based Sobel edge detection algorithm produced an edge map then a thinning algorithm then reduced strong border edge lines so they could be excluded. Next, the edge areas were measured, and this value was divided by the cross-sectional area of the nucleus.

### Quantification of immunofluorescence images

For quantification of CD73 and Runx2 protein fluorescence, 20x images of cells stained for DAPI, actin, CD73, and Runx2 were imaged together. ImageJ was used for measuring the intensity values for each channel. Briefly, in the Runx2 channel, cells were randomly chosen throughout the well, and the cell and nucleus shape were outlined and added as a region of interest (ROI). The mean intensity was then calculated by Image J for the nucleus in the Runx2 channel and the cell shape ROI was added to the CD73 channel where that mean intensity was measured. Then, the nucleus image was were subtracted from the cell image to obtain an ROI of the cell excluding the nucleus area. This new ROI was opened in the Runx2 channel where the cytoplasmic Runx2 mean intensity was calculated. The Runx2 mean intensity within the nucleus and the cytoplasm respectively were divided to obtain a ratio of nuclear Runx2 over cytoplasmic Runx2 mean intensity.

### Quantification of heterogeneity

Heterogeneity of the quantified parameters such as cell and nuclear geometrical phenotype was evaluated using a common measure of dispersion, the interquartile range (IQR) (quartile 3 - quartile 1). IQR as a measure of dispersion can be beneficial for quantifying the biological data specifically due to the irregularity that can occur between biological replicates. The usual drawback for IQR is that only a subset of the data points is accounted for. However, performing regression deletion diagnostics revealed that each linear model, even after a log transform, contains several influential outliers within our datasets. Using IQR as a measure of data dispersion reduces the effects of these extreme values [33]. Concurrently, a large consideration for other measures of dispersion is the distribution of the data, with a normal distribution being a prerequisite for proper representation. Visualization of the density plots of the data (Figure S2-5) revealed that our data has a variety of skewed distributions, however, IQR is not tied to the symmetry or distribution of the data, unlike standard deviation which would lose its effectiveness [34].

### Cell treatments with cytoskeleton modifying drugs

To understand the role of mechanical integrity of the cytoskeleton on the chromatin architecture and the nuclear phenotype, cytochalasin D (ThermoFisher Scientific) at a concentration of 0.02 μM or 0.05 μM and Y27632 (Tocris Biosciences) at concentrations of 2 μM or 10 μM were applied to P5 cells 24 hours after seeding. All the control groups for these experiments were vehicle treated (DMSO). Cells were then fixed after 2 days of treatment, for subsequent staining and imaging. The combination drug treatments combine the two lowest concentrations of Cytochalasin D (0.02 μM) and Y27632 (2 μM) together.

### Statistical Analysis

Statistical analysis was performed using Student’s t-test (RStudio [35, 36]). Results are expressed as mean ± standard deviation, with differences considered statistically significant at p < 0.05 or p < 0.01, as specified in the figure legends. Sample and replicate numbers are provided in the figure legends.

## RESULTS

### Soft substrates promote a more homogeneous population of cell phenotype over passaging

MSC phenotype is known to be affected by the substrate mechanical stiffness. We investigated how the quantitative measures of the cell phenotype and their heterogeneity is affected by a wide range of stiffness representing the physiologically healthy bone marrow (soft), pathological fibrotic tissue (medium) and tissue culture substrate (stiff) (Figure 1, Figure S2A). Passage 3 (P3) MSC were cultured to 90% confluency and subcultured to passage 4 (P4) to quantify how the mechanics of the substrate affects the heterogeneity in the cell area, and the F-actin intensity over one passaging. It was observed that on stiff substrates, the cell area was higher, and the cells were flatter, whereas the cells grown on the medium and soft substrates better maintained a smaller cell area and spindle-like shape, characteristics of MSC (Figure 1A). Quantification of these observations revealed that there was significant spreading on the stiff substrate over one passage, with the cells on the soft and medium substrates remaining significantly smaller than the cells on the stiff substrate even after one passage (Figure 1 B, E). As an indicator of MSC population heterogeneity, the interquartile range (IQR) of the area measurements was calculated (Figure 1C). The IQR values indicated that the soft and medium substrate significantly reduced the variation of cell area over one passage when compared to cells grown on stiff substrates (Figure 1 C, E). The IQR values further revealed that the medium substrate homogenized the cell area over one passage, while the stiff substrate cell area became more heterogeneous, whereas on soft substrate the heterogeneity remained low over one passage. Another observation from high resolution imaging was the increasing intensity of the phalloidin (F-actin) stain over passaging on stiff substrates (Figure 1A, *top right inset image*). The intensity ratio of the F-actin significantly increased from P3 to P4 in the stiff and medium groups, but significantly decreased on the soft substrate (Figure 1D, E). Collectively, the IQR of cell area, and the relative F-actin intensity indicates that MSC phenotype was less heterogeneous on softer substrate over one passaging.

### Soft substrates promote a more homogeneous nuclear phenotype over passaging

After the investigation of cell area and cytoskeleton structure, we examined the effect of substrate stiffness on the nucleus phenotype over passaging (Figure 2, Figure S2B). Differences in size and shape were observed from P3 to P4 on the various substrates (Figure 2A). Upon quantification of nuclear area and circularity (Figure 2B, F), we found that the nuclei for cells on the medium and stiff substrates became significantly larger in area and more rounded (circularity values closer to 1) after one passage thus indicating a deviation from MSC phenotype. On the soft substrate in P3, nuclei were significantly smaller and more elongated, which was maintained in P4 (Figure 2B, F). Subsequently, the IQR of nucleus area and circularity was examined, again, as an indicator of the heterogeneity of the MSC population phenotype at the nuclear level. IQR values increased for nucleus area and circularity on the medium and stiff substrate but decreased on the soft substrate (Figure 2D, F) over one passage. Therefore, the MSC nuclear phenotype was maintained on soft substrate over one passage.

**Figure 1.**
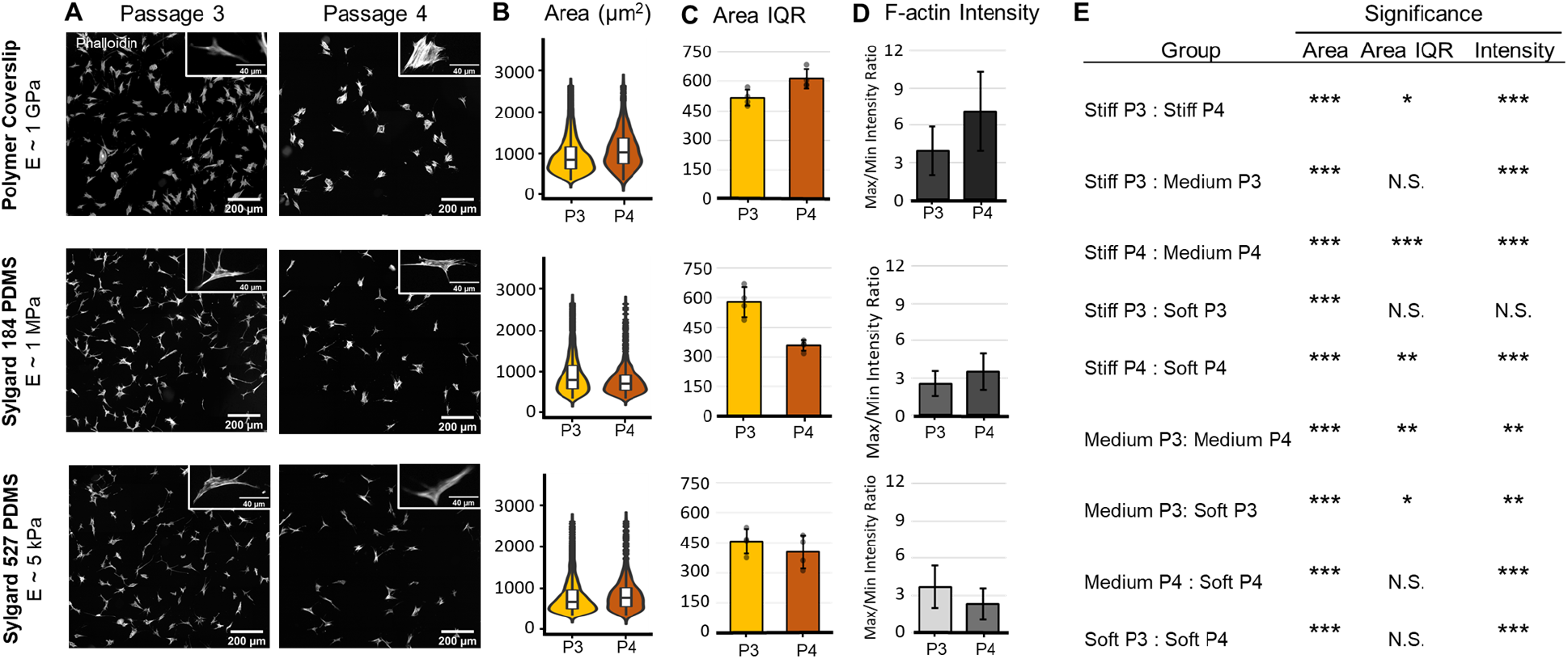
Effect of substrate stiffness on human BM-MSC cytoskeleton phenotype over one passage. (A) Representative images of F-actin staining over one passage on substrates with varying stiffnesses. (B) Violin plots to show complete distribution of the data with embedded boxplot (n = 1000 cells) for cell area. (C) Quantification of the interquartile range (IQR) of the parameter cell area from 4 technical replicates to represent heterogeneity of the MSC population. (D) F-actin intensity quantification of MSCs cultured on PDMS substrates in passage 3 (P3) and passage 4 (P4) (n ∼ 30 cells). (E) Table of p-values for various comparisons in each quantity. Stiff = 1 GPa, medium = 1 MPa, soft = 5 kPa. Error bars represent ± SD. N.S. (Not Significant): p>0.05, *p<0.05, **p<0.01, ***p<0.001.

**Figure 2.**
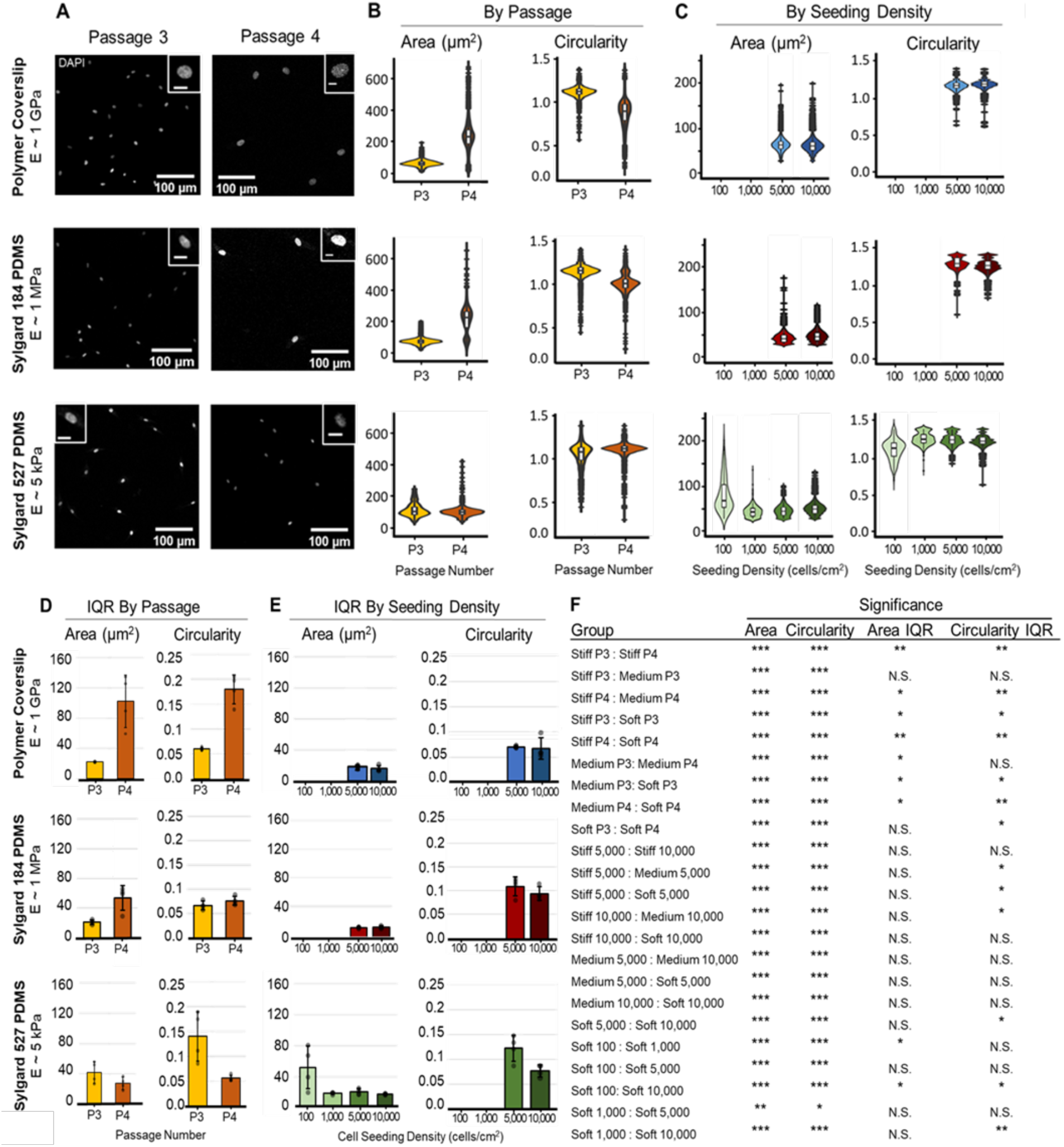
Effect of substrate stiffness and cell seeding density on human BM-MSC nucleus phenotype. (A) Representative images of DAPI stained MSCs over a single passage seeded at 5,000 cells/cm^2^. Inset scale bar: 10 μm. (B) Nucleus area and circularity quantification over one passage seeded at 5,000 cells/cm^2^, visualized with violin plots to show the complete distribution of the data with embedded boxplot (n = 1000 cells). Circularity is a measure of elongation with 1 representing a perfect circle and an increasing deviation from 1 represents a more elongated shape. (C) Nucleus area and circularity quantification in passage 4 cells with varied seeding densities (n = 1000). (D,E) Quantification of the interquartile range (IQR) of the nucleus area and circularity based on 4 technical replicates to represent heterogeneity of the MSC population. (F) Table of comparisons and the corresponding p-value. Error bars represent ± SD. N.S. (Not Significant): p>0.05, *p<0.05, **p<0.01, ***p<0.001.

### Seeding density has little impact on nucleus phenotype, but significantly affects chromatin organization

After determining the effect of substrate stiffness and passaging on MSC cell and nuclear phenotype, we wanted to examine their influence in conjunction with seeding density to explore the effect of substrate mediated cell communication on the nucleus phenotype (Figure 2, S3). P4 cells were seeded on the stiff, medium, or soft substrates at increasing seeding densities and the nucleus phenotype was analyzed. Cells grown at 5,000 and 10,000 cells/cm^2^ had the smallest area on the soft substrate and were more elongated on the soft and medium substrates compared to the stiff substrate (Figure 2C, F). The IQR values at these higher seeding densities for nucleus area and circularity were largely unaffected by differences in seeding density. A wider array of seeding densities was tested on the soft substrate to explore the limits of cell communication through the compliant substrate. At the sparse seeding density of 100 cells/cm^2^, the shape of the nucleus varied widely in area and circularity (Figure 2C). This was corroborated in the significantly large IQR values for this low seeding density, representing higher phenotypic heterogeneity (Figure 2E, F). Taken together, these results indicate the substrate stiffness and not the seeding density is the most influential factor in maintaining a homogeneous MSC population.

Chromatin architecture and organization is closely correlated to the MSC phenotype and function, as demonstrated in previous studies [37, 38]. To understand the effect of substrate stiffness and cell seeding density on the chromatin organization, we quantified the percentage of chromatin segregation. A larger percentage of segregated chromatin represents an open chromatin architecture – a hallmark of MSC. As the substrate became softer, the chromatin became significantly more segregated (Figure 3A), with a similar significant effect seen as the seeding density became higher. Of note, the stiff substrate as well as low seeding densities on soft substrate resulted in a diffused chromatin architecture with low chromatin segregation (Figure 3C). Concurrently, examining the IQR values of the chromatin segregation index showed a foreseeable pattern, that more chromatin segregation led to a larger IQR value (Figure 3B), representing a population of MSC with higher differentiation potential. These findings show that cell communication along with a higher seeding density through a soft substrate leads to a significantly open chromatin architecture.

**Figure 3.**
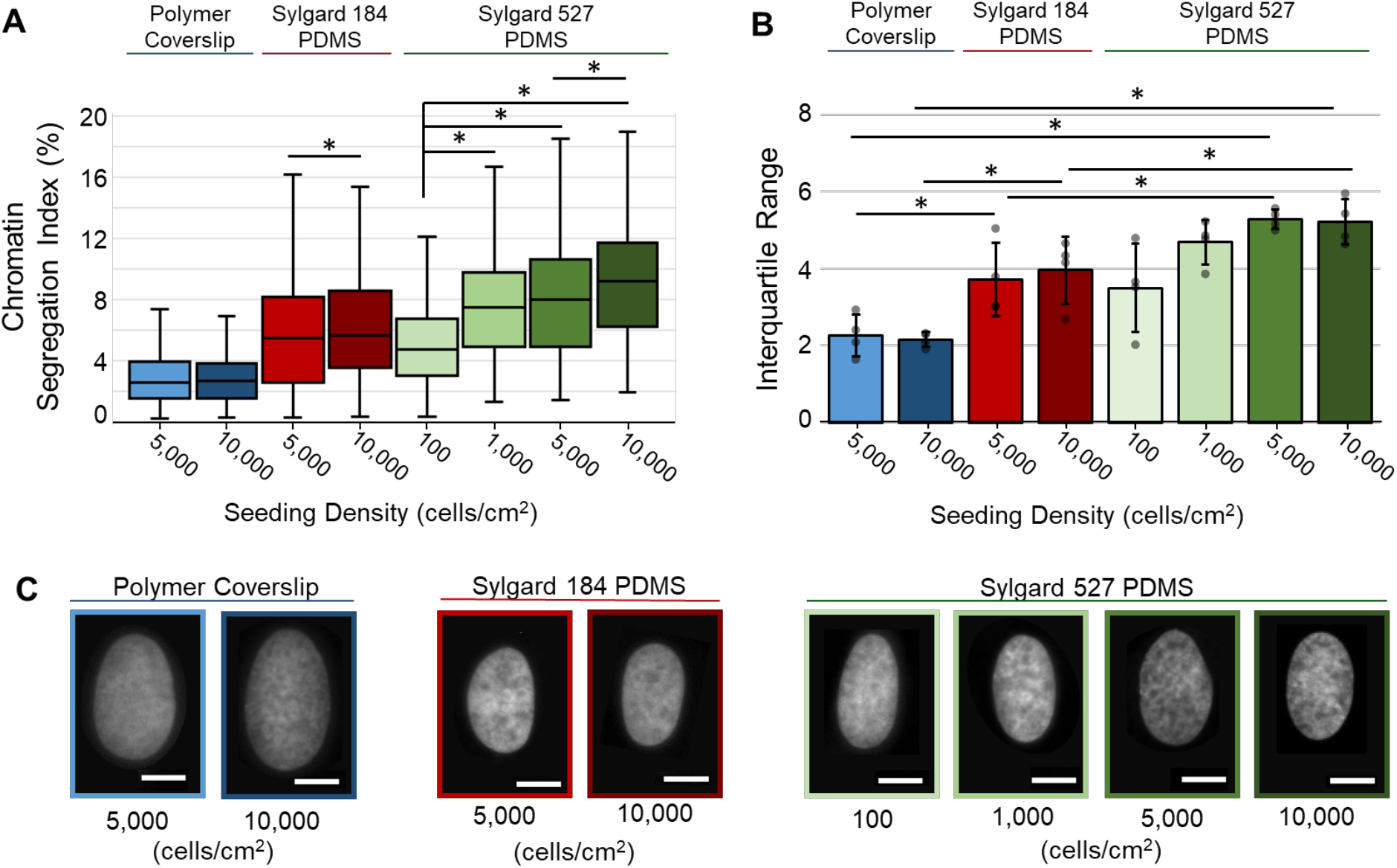
Effect of cell seeding density in combination with substrate stiffness on the chromatin architecture of hMSCs. (A) Quantification of the percent chromatin segregation in passage 4 cells (n = 40). (B) Quantification of the interquartile range (IQR) of the percent chromatin segregation from 4 technical replicates to represent heterogeneity of the MSC population. (C) Representative images of DAPI stained MSC nuclei. Scale bar:10 μm. Error bars represent ± SD. *p<0.05, ***p<0.001.

### Functional indications of substrate mechanics driven heterogeneity

To examine the functionality of MSCs on the varying substrates in more detail, Passage 4 MSCs were stained for the MSC-specific marker, CD73 and the osteogenesis precursor protein, Runx2. We found a strong relationship between the level of CD73 and level of Runx2 on the stiff and soft substrates, but no relationship on the medium stiffness substrate (Figure 4A). On soft substrates, cells showed a higher CD73 intensity and lower localization of Runx2 to the nucleus. Stiff substrates however, showed low CD73 intensity with a high ratio of Runx2 in the nucleus. Localization of Runx2 to the nucleus is an indicator that the cells begin to commit to an osteoprogenitor lineage. Soft substrates showed significantly higher CD73 intensity (Figure 4B, S4A) and significantly lower Runx2 nuclear localization (Figure 4D, S4B) compared to the stiff and medium substrates. When examining the heterogeneity of CD73, we saw a significant increase in the medium and soft substrates compared to the stiff substrate (Figure 4C), similar the trend of chromatin segregation heterogeneity. CD73 is specific to MSCs, which exhibit differentiation potential and this high IQR value represents various lineage opportunities. There were no significant differences in IQR for the Runx2 nucleus localization (Figure 4E). Overall, we found that soft substrate has higher population of MSC-marked cells with Runx2 that is less localized to the nucleus than cells on stiff and medium stiffness substrates.

**Figure 4.**
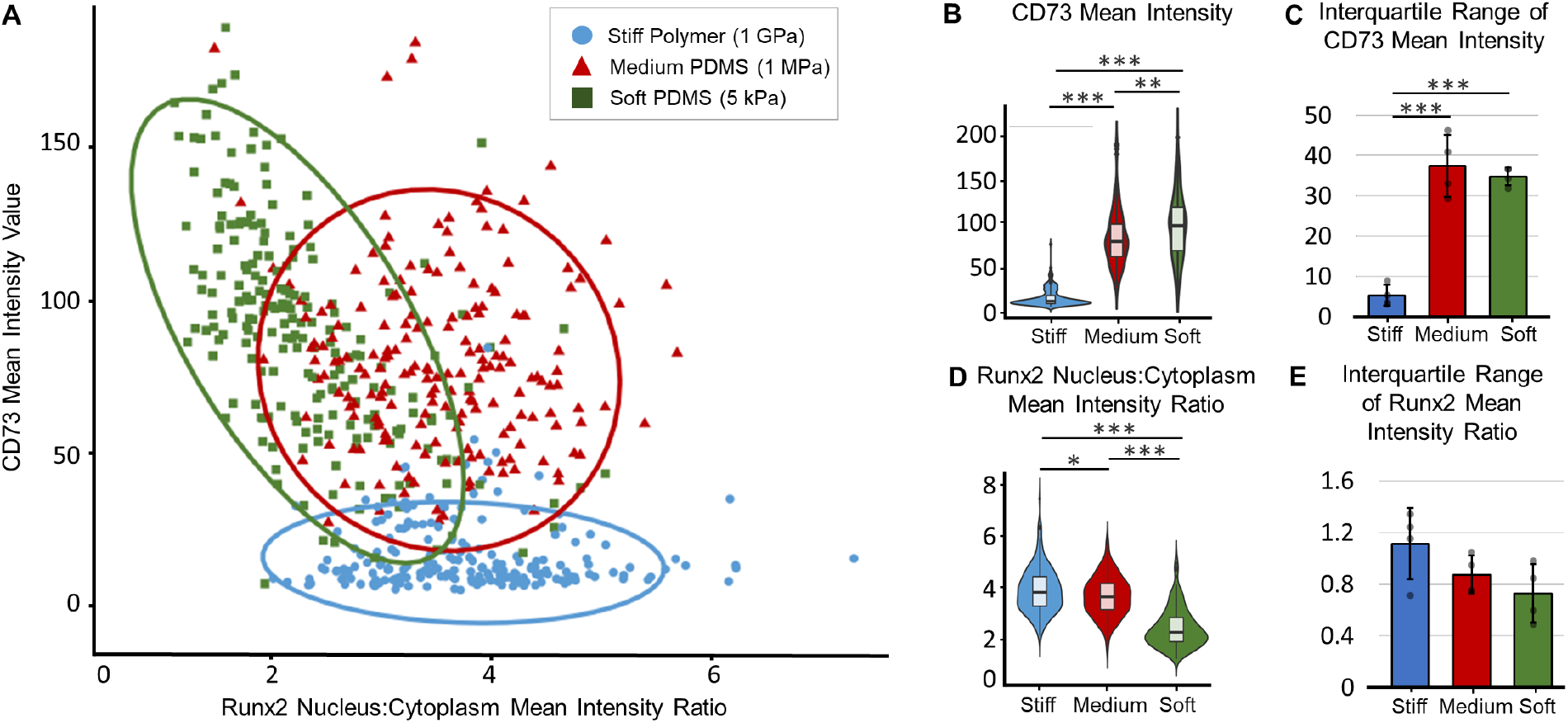
Mean fluorescent intensity analysis and heterogeneity quantification of P4 MSCs seeded at 5,000 cells/cm^2^. (A) Scatter plot of CD73 mean intensity and Runx2 nucleus to cytoplasm mean intensity ratio for cells cultured on varied substrate stiffnesses. Each data point represents one cell. 95% confidence level ellipses for a multivariate t-distribution were included as a visual indicator of correlation. (B) Mean intensity values of CD73 visualized with violin plots to show full distribution of the data with embedded boxplot (n = 200 cells). (C) Quantification of the interquartile range (IQR) of the CD73 mean intensity values from 4 technical replicates to represent heterogeneity of the MSC population. (D) Ratio of the mean intensity value of Runx2 in the nucleus divided by the mean intensity value of Runx2 in the cytoplasm (n = 200 cells). (E) Quantification of the interquartile range (IQR) of the Runx2 nucleus to cytoplasm mean intensity ratio from 4 technical replicates to represent heterogeneity of the MSC population. Error bars represent ± SD. *p<0.05, **p<0.01, ***p<0.001.

### Cytoskeletal tension significantly affects cell and nuclear phenotype but not the heterogeneity

Now that we have determined that substrate stiffness plays a significant role on the MSC phenotype and the level of phenotype heterogeneity, we attempted to isolate the effects of cytoskeletal tension from cell-cell mechanical communication through the substrate (Figure 5, S5). Passage 5 (P5) cells on stiff substrates were treated either with cytochalasin D (cyto-D), Y27632 or a combination of both drugs. Treatment with cyto-D inhibits the actin polymerization, and treatment with Y27632 inhibits the Rho-associated protein kinase (ROCK), impeding actomyosin contraction. The ROCK inhibition led to smaller nuclear area, increased elongated nuclear shape and the overall smaller cell area (Figure 5A, C), thus recovering the MSC phenotype. However, the analysis of the IQR values, as an indicator of MSC population heterogeneity, revealed that cytoskeletal mechanics inhibitors had no significant effects on IQR (Figure 5B, D), at least in the late passage cells on a stiff substrate. Lastly, we wanted to examine if those cytoskeletal mechanics inhibitors affect the formation of stress fibers. There were no significant changes in F-actin intensity ratio (Figure 5E) compared to the control group. There results indicate that cytoskeletal tension indeed affects the MSC phenotype but not their phenotypical heterogeneity.

**Figure 5.**
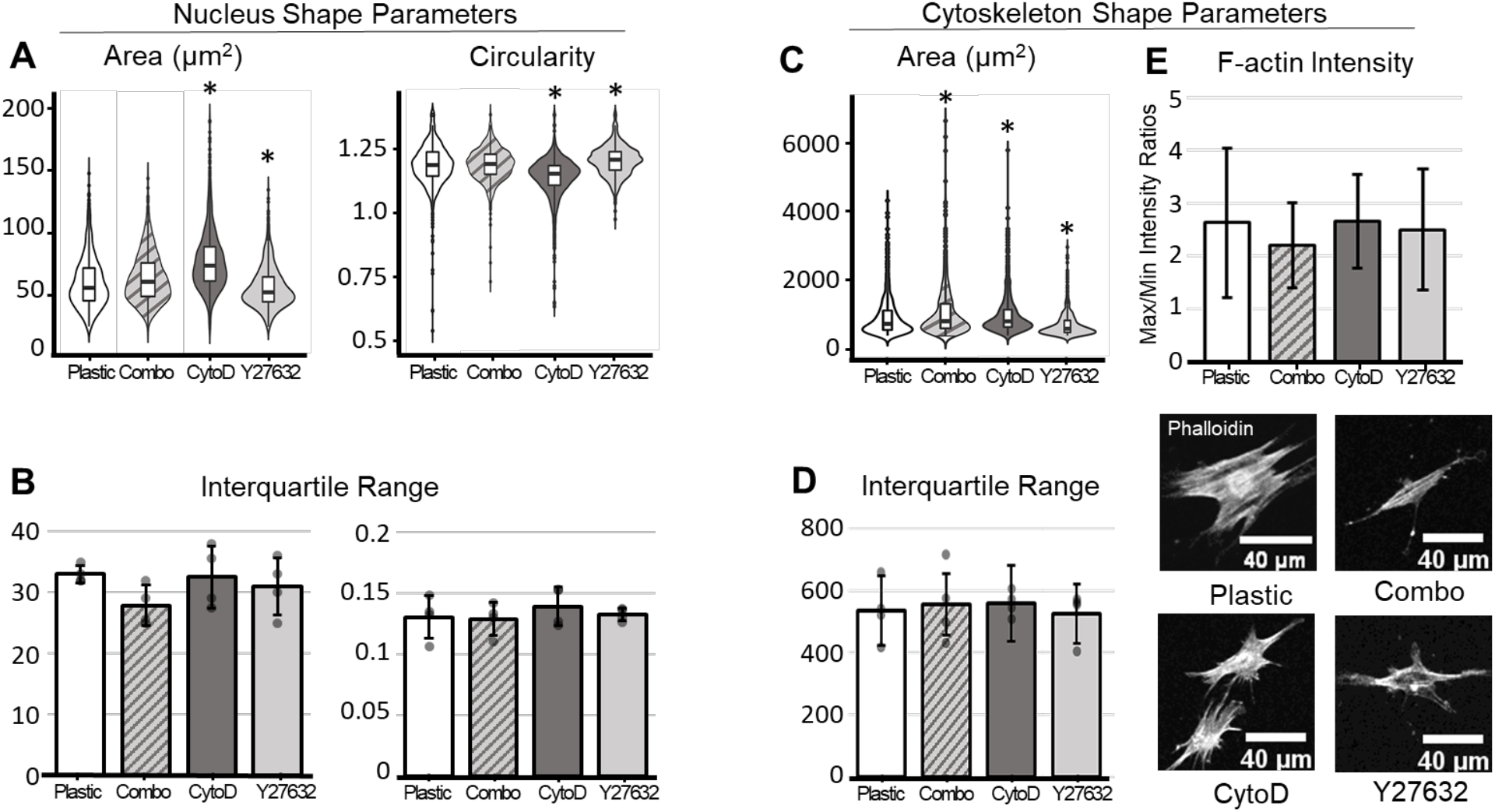
Effect of mechanics inhibitors on the phenotype of hMSCs. Passage 5 cells were cultured on a stiff substrate and were treated with cytochalasin D (cyto-D), the ROCK inhibitor Y27632, or a combination treatment of cyto-D and Y27632. (A) Nucleus area and circularity quantification visualized through violin plots to show the distribution of the data with embedded boxplots (n = 1000 cells). (B) Quantification of the interquartile range (IQR) of the nucleus area and circularity from 4 technical replicates to represent heterogeneity of the MSC population. (C) Cell area and quantification visualized through violin plots to show the distribution of the data with embedded boxplots (n = 1000 cells). (D) Quantification of the interquartile range (IQR) of the cell area from 4 technical replicates to represent heterogeneity of the MSC population. (E) F-actin intensity quantification of hMSCs cultured on tissue culture plastic in passage 5 with various mechanics inhibitor treatments (n = ∼30 cells). Representative images of the phalloidin stained cells from each treatment group. Error bar: SD. *p<0.05.

### Cytoskeletal tension is reinforced by the perinuclear actin cap

The increased cytoskeletal tension in cells on stiff substrates was also associated with an observed dome of actin filaments at the apical surface of the MSC nucleus, known as the perinuclear actin cap (Figure 6A, B). Quantification of the proportion of cells with perinuclear actin caps revealed a relationship between substrate stiffness and seeding density and the perinuclear actin formation. Presence of perinuclear actin caps was more frequent in cells at lower cell density and on the stiff substrate, with instances decreasing significantly for cells on the soft substrate and at a higher seeding density (Figure 6C, D). This corresponds with the increased cell and nucleus area and F-actin intensity on stiff substrates quantified earlier (Figure 1B, C and Figure 2B, C). Together, these results suggest that the perinuclear actin cap formation is involved with cytoskeletal tension mediated stress fiber formation around the nucleus.

**Figure 6.**
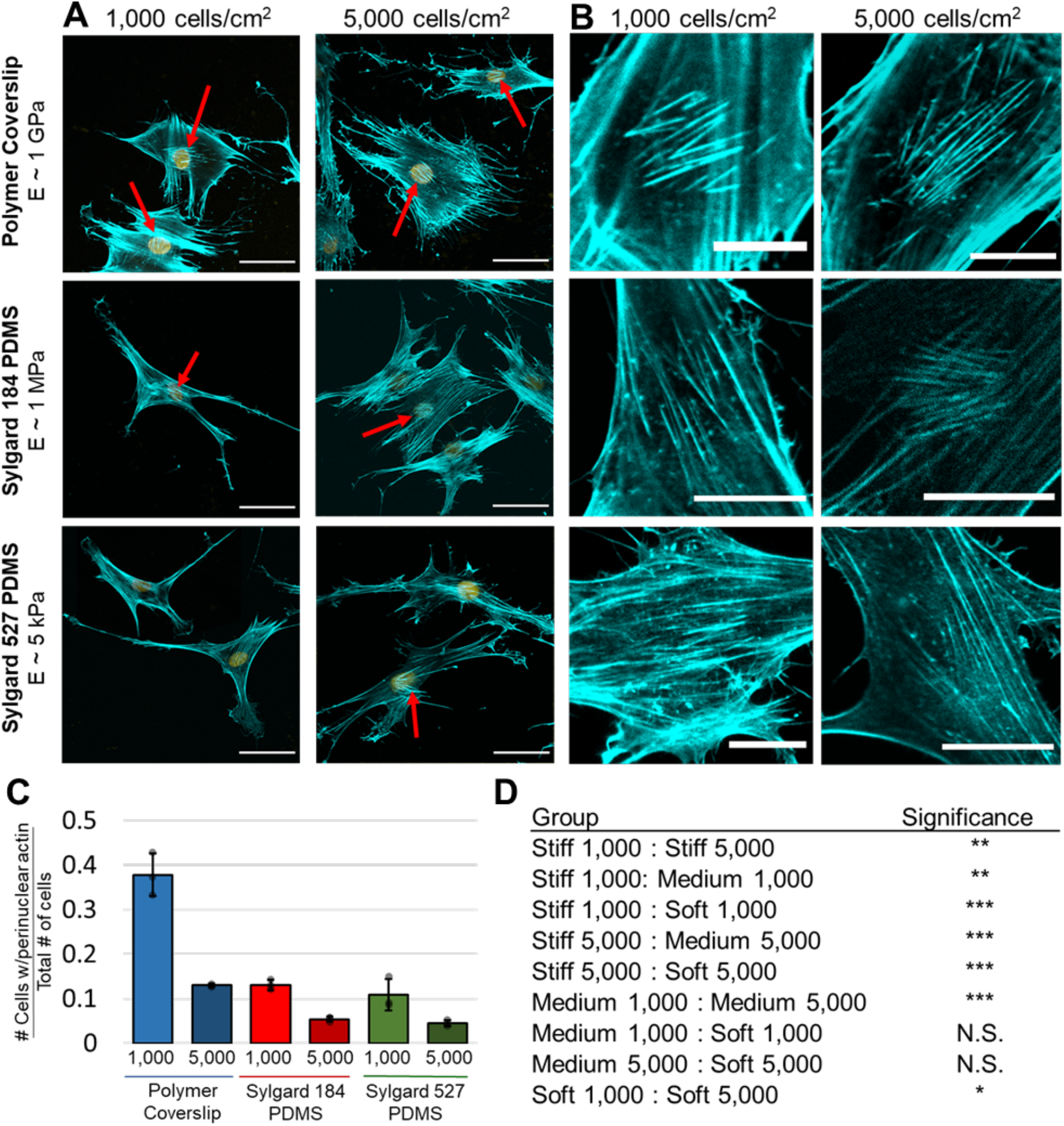
Effect of substrate stiffness and cell seeding density on perinuclear actin formation in BM-MSCs. (A) It was observed that MSCs displayed less perinuclear actin on soft substrates. Red arrows point to cells with visible perinuclear actin. Scale bar: 50 μm. (B) High resolution imaging of identified perinuclear actin in MSCs shows a clear reduction in perinuclear actin on soft substrates. Scale bar: 20 μm. (C) Ratios were quantified from 3 technical replicates to examine the number of cells that displayed perinuclear actin compared to the total amount of cells in the image (n = 150 for 1,000 cells/cm^2^ group, n = 650 for 5,000 cells/cm^2^ group). (D) Table of p-values for various comparisons in each quantity. Stiff = 1 GPa, medium = 1 MPa, soft = 5 kPa. Error bars represent ± SD. N.S. Not Significant: p>0.05, *p<0.05, **p<0.01, ***p<0.001.

## DISCUSSION

In this study, we provided evidence of phenotypical heterogeneity of MSC during 2D expansion culture. The heterogeneity considered throughout this study was evaluated using a common measure of dispersion, the interquartile range (IQR). Mechanistic insight into the source of substrate mechanics-driven heterogeneity, further influenced by the substrate-mediated cell-cell communication was examined. We found that substrate stiffness plays a key role in the heterogeneity of the MSC phenotype. Soft substrates (∼5 kPa) promoted a more homogeneous population of cells with the characteristic MSC cytoskeleton and nuclear phenotype, accompanied by increased expression of CD73, which is associated with a high degree of stemness [39]. Conversely, MSCs grown on the stiff substrate (∼1 GPa) not only showed phenotype abnormalities, presence of perinuclear actin caps, and Runx2 nuclear localization, in line with existing reports [40–42], but also, they displayed a heterogeneity of phenotype at the population level. These findings are associated with increased cellular senescence [31], decreased proliferation [40], and decreased differentiation potential [12] in MSCs.

It is well-known that MSC are exposed to biochemical and biophysical signals that can determine lineage commitment [43]. Differentiation via biochemical cues have been extensively studied, while mechanisms of cellular mechanotransduction are becoming increasingly understood. The combination of cell-generated forces and matrix mechanics to coordinate the MSC population needs further investigation to understand these specific interactions. We hypothesized that neighboring cells would communicate through the soft substrate to promote a homogeneous population of MSCs. However, we found that seeding density had little impact on the heterogeneity of nuclear shape but had a significant effect on the chromatin organization. In the extremely low seeding density case at 100 cells/cm^2^, the cells were very far apart, impeding mechanical signaling, resulting in an extremely varied phenotype, spanning from fibroblast like flat phenotype to MSC like spindle phenotype with a closed chromatin architecture. An open chromatin architecture, as seen in cells on the softer substrate, is associated with transcriptional activation and is a hallmark of stem cells [44]. These changes in chromatin organization are most likely driven by the mechanical communication to neighboring cells through the substrate because actin fiber intensity was consistent between each seeding density. Accordingly, the increased IQR of chromatin segregation when cells were grown on soft substrates, supports the ability of MSCs to be in varying transcriptional states.

The phenotypic trends towards the MSC-like phenotype and chromatin segregation patterns on soft substrates was substantiated upon CD73 immunofluorescence. CD73 enrichment indicates MSCs and the heterogeneous enrichment patterns is associated with a reparative properties and anti-inflammatory activity [45]. The nuclear to cytoplasmic ratio of Runx2 was analyzed as an indication of early indication of osteogenic differentiation. When Runx2 is translocated into the nucleus, it induces transcriptional expression of osteogenesis-related genes [46]. The pattern that emerged due to the high nuclear Runx2 in cells on the stiff substrate is consistent with the phenotype of cells on the stiff substrate.

A potential approach to alter the heterogeneity dynamics of a population of MSCs during in vitro culture was to disrupt the cytoskeleton tension to impose a lower tensile stress inside the cells. Disrupting the cytoskeletal tension with the ROCK inhibitor Y27632 resulted in cells with a significantly restored MSC cell and nucleus phenotype. Cell area was lowered, nucleus became smaller in area and more elongated, representing the MSC phenotype. A contributing factor to this recovered phenotype after Y27632 treatment is the disruption of the LINC complexes connecting the nucleus to this actin cap, decreasing cellular tension [40]. ROCK inhibition is also known to prevent osteogenic differentiation of MSCs [26]. The phenotype heterogeneity, however, was not significantly affected by inhibiting actin polymerization or actomyosin contraction. Studies have been performed to understand the effects of these cytoskeleton modifying drugs on MSC functionality, but the effects on the population heterogeneity is not specifically explored. Our findings from this study suggests that stiffness-dependent heterogeneity does not rely on actin formation and actomyosin contractions as we hypothesized, and we explain the observation as follows. Cytoplasmic dynamics are faster than the intranuclear dynamics. Therefore, during continuous culture to passage 5 on stiff substrate, the amount of existing F-actin fibers that developed might have outweighed the effects of Y27632, likely leading to irreversible heterogeneity. Overall, this finding might be attributed to the lower plasticity of MSC at later passage.

Recognizing the natural heterogeneity dynamics of a cell population and understanding its source allows for minimization of adverse outcomes in efficacy or safety from one patient to the next. This study provides potential targets in the intracellular mechanobiological pathways to control the heterogeneity dynamics of MSCs during in vitro expansion. Future studies would involve single cell RNA-sequencing to further quantify the heterogeneity in MSC phenotype and explore the possible pathways to understand the mechanistic origins of heterogeneity.

## ACKNOWLEDGEMENTS

The authors would like to thank the funding from multiple units at Colorado State University. The authors are grateful to Jack Forman for assistance with image processing.

## AUTHOR CONTRIBUTIONS

Conceptualization, S.K. and S.G.; Experiments and methodology, S.K. and S.G.; Data analysis and statistics, S.K. and Z.A.; Writing, S.K. and S.G.; Resources and funding acquisition, S.G.

## SUPPORTING INFORMATION

### Supplementary Figures

**Supplementary Figure 1.**
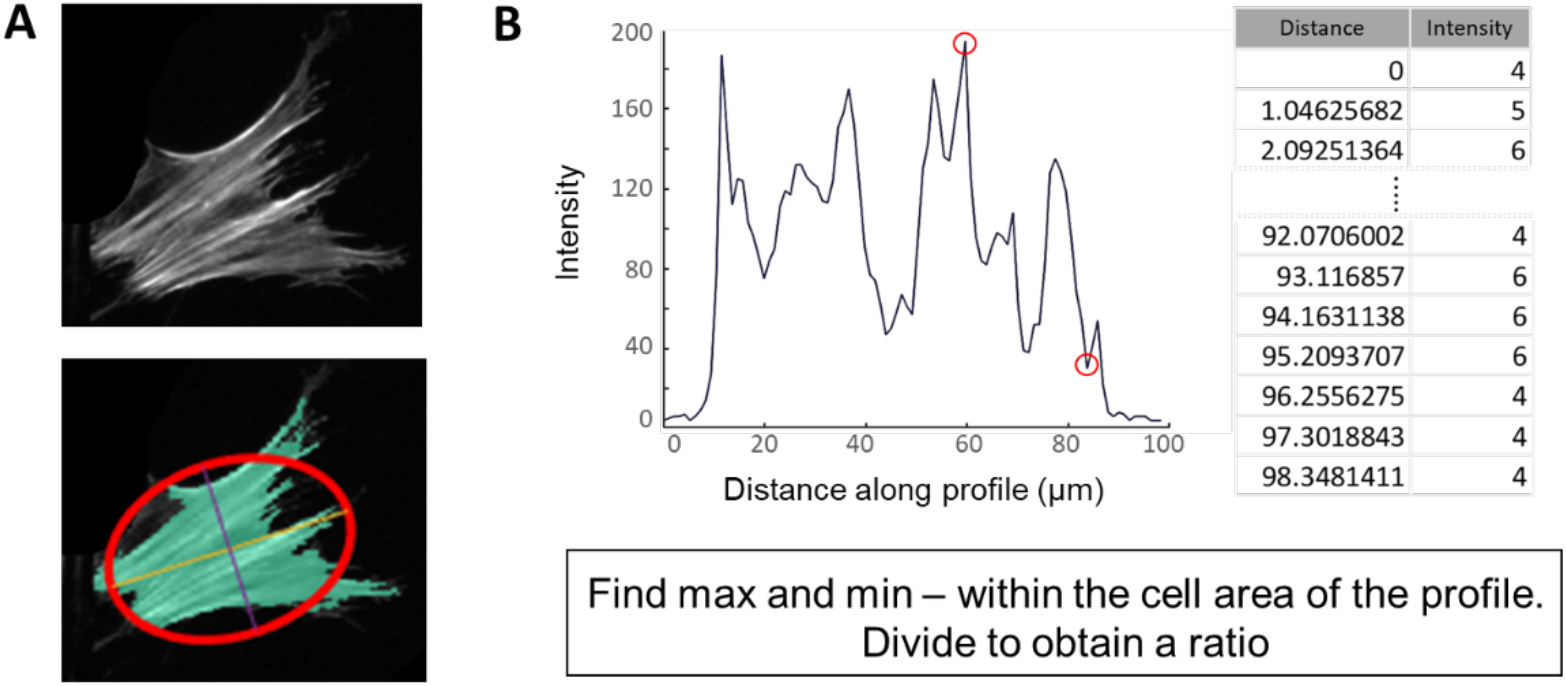
F-actin intensity quantification method. A custom MATLAB code was used to identify individual cells in an image (colored for confirmation). An ellipse fit based on the major and minor axes. Intensity maps and corresponding tables are then printed for each major and minor axis to the screen and saved. The cell number is noted on the graph and in the tables for reference. In general, the minor axis is used because it is usually aligned perpendicular to the actin fibers, but this is confirmed for each cell. Then the maximum value was divided by the minimum value within the bounds of the cell area to obtain an intensity ratio.

**Supplementary Figure 2.**
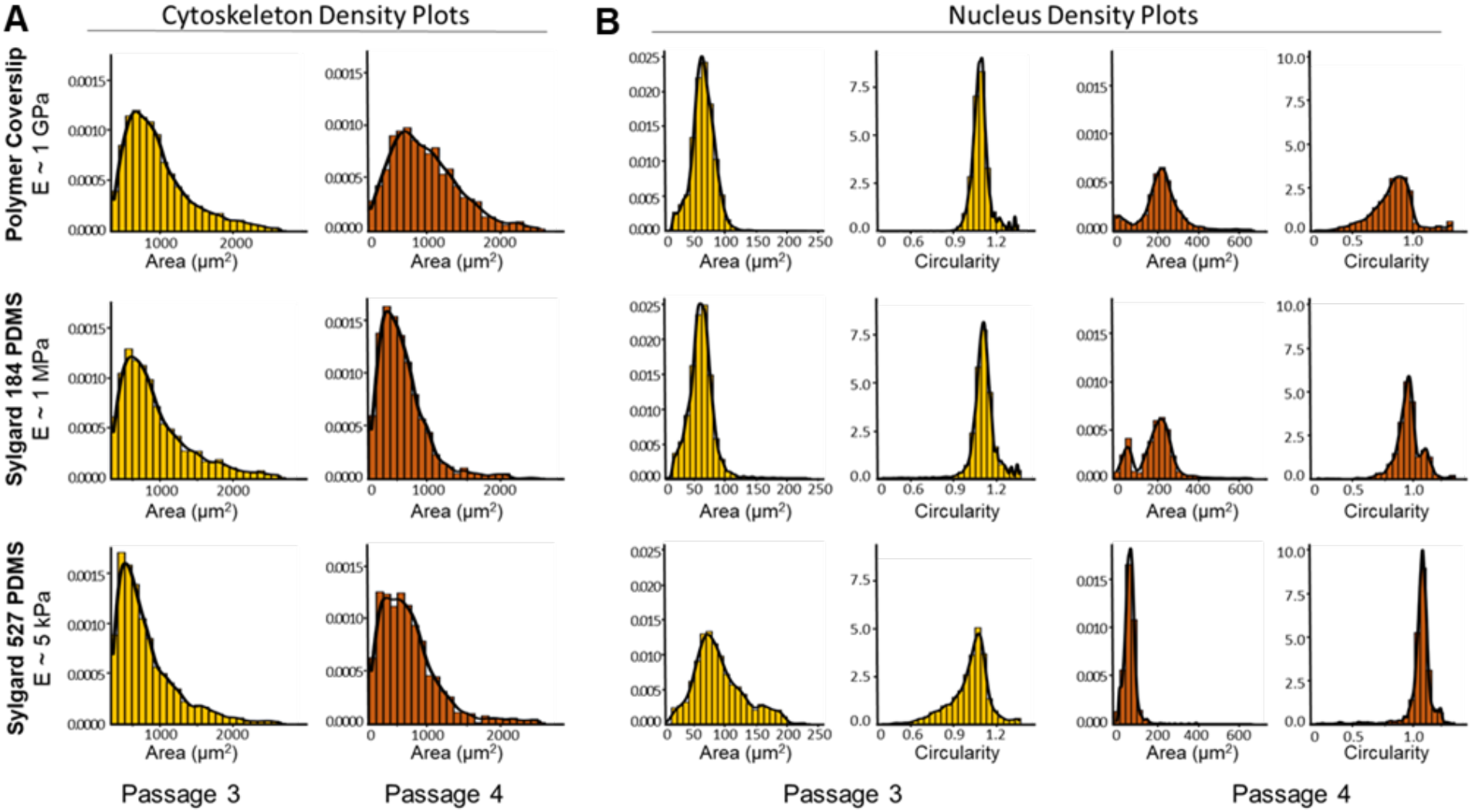
Density plots using a kernel density estimate to show the probability density function of each variable in Passage 3 or Passage 4. (A) Cytoskeleton area quantification density plots for the violin and box plot reported in Figure 1B. (B) Nucleus area and circularity quantification density plots for the violin and box plots reported in Figure 2B.

**Supplementary Figure 3.**
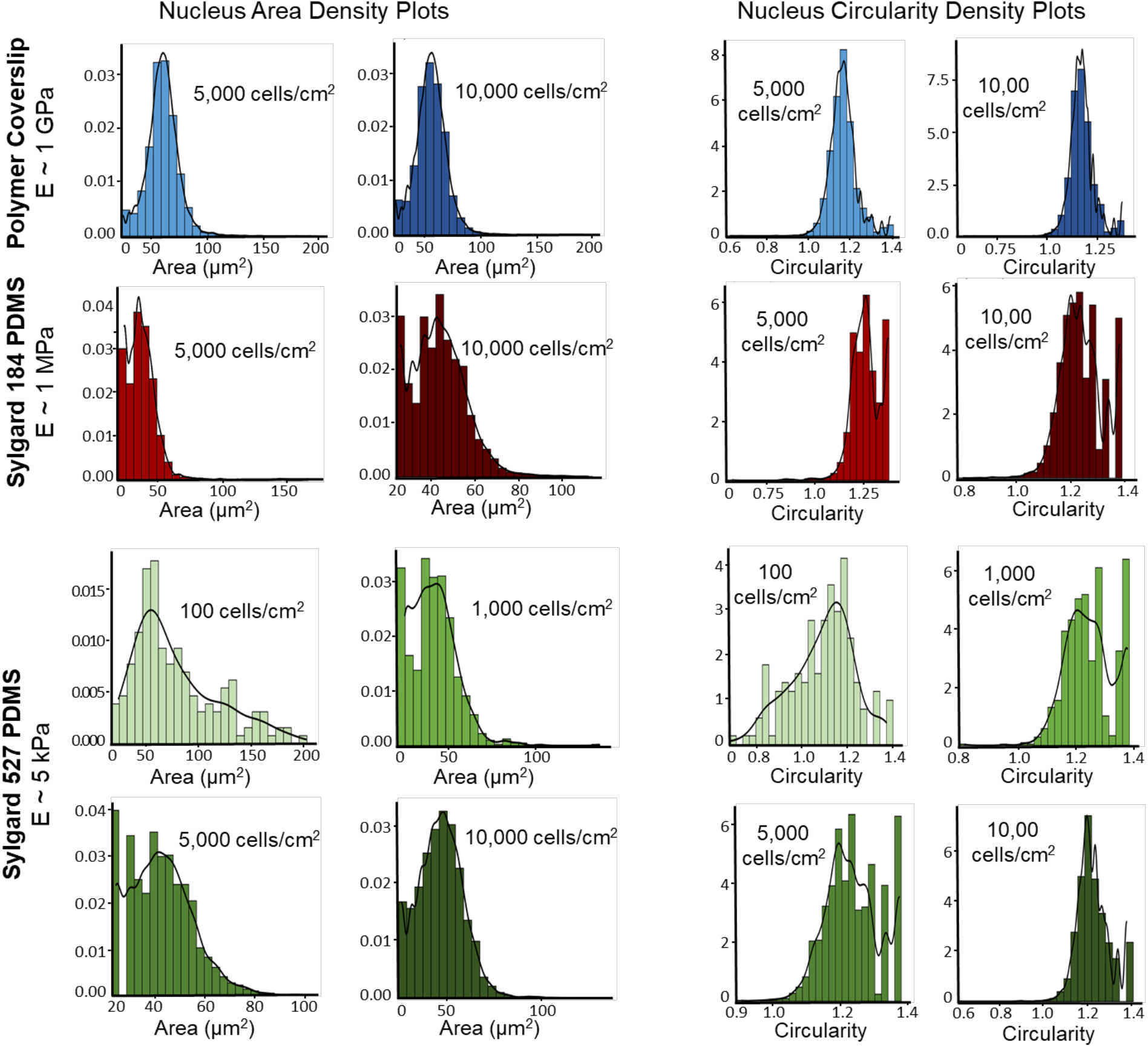
Density plots using a kernel density estimate to show the probability density function of each variable under various seeding densities. Nucleus area and circularity were quantified on each substrate stiffness and under increasing seeding densities, listed in each graph. These density plots correspond to the violin and box plots reported in Figure 2C.

**Supplementary Figure 4.**
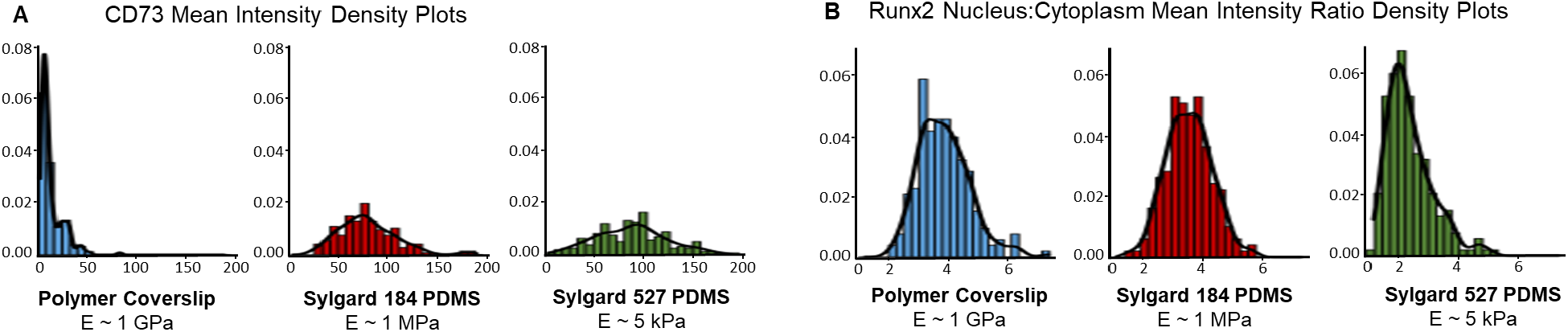
Density plots using a kernel density estimate to show the probability density function of (A) CD73 mean fluorescent intensities or (B) Runx2 nucleus to cytoplasmic mean intensity ratios on each substrate stiffness. These density plots correspond to the violin and box plots reported in Figure 4.

**Supplementary Figure 5.**
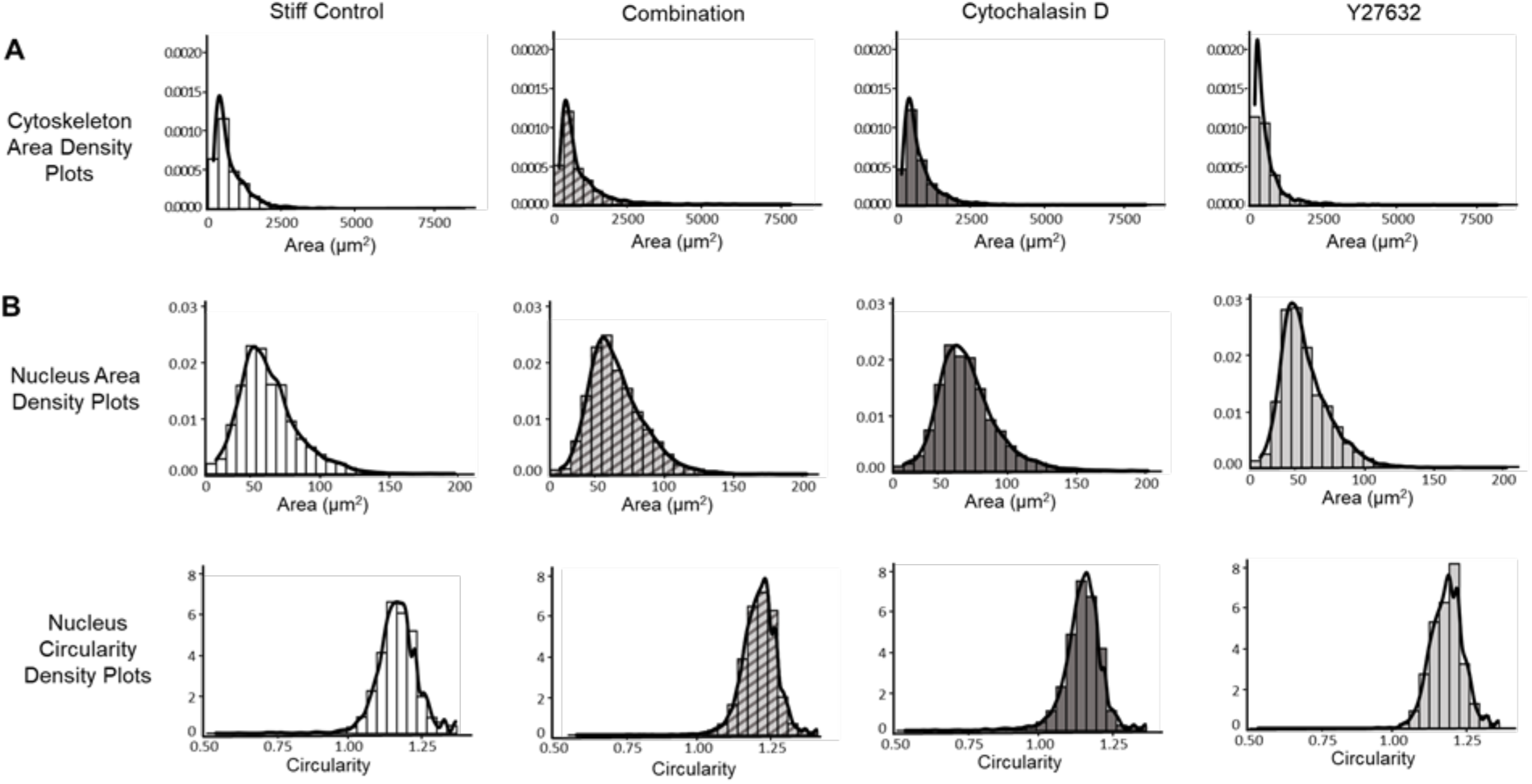
Density plots using a kernel density estimate to show the probability density function of each variable after undergoing treatment with various drug treatments. Passage 5 cells were cultured on a stiff substrate and were treated with cytochalasin D (cyto-D), the ROCK inhibitor Y27632, or a combination treatment of cyto-D and Y27632. (A) Cytoskeleton area quantification density plots for the violin and box plot reported in Figure 5C. (B) Nucleus area and circularity quantification density plots for the violin and box plots reported in Figure 5A.

### Quantification of stress fiber formation

Stress fiber intensity quantification was performed using a custom MATLAB code. Cropped confocal images were converted to grayscale and a pixel value threshold was set to create a binary mask around that threshold. Then arbitrary structuring elements were created through the ‘strel’ function, which created masks over each identified cell in the image. The edges of the identified elements were smoothed and small holes in the element was filled so a solid mask could be visualized over the original image. To avoid any background noise that may have been included, a size filter was run to exclude any elements that would be too small to be considered a cell. The final allowable elements were labeled for later reference and then measured with ‘regionprops’, which included an estimated ellipse from the major and minor axes of each identified cell. Then the intensity maps and corresponding tables were separately printed and saved for the major and minor axis of each numbered cell. Generally, the minor axis aligned perpendicular to the actin fibers, though each cell was manually checked to confirm. The code relied heavily on aligned F-actin fibers so in some experiment groups with harder to discern F-actin fiber alignment, such as the Y27632 treated cells, the axis of intensity calculation was done manually in ImageJ. After the perpendicular axis was determined and located in the output table for each identified cell, the maximum and local minimum, was then divided to obtain a ratio of intensities (Figure S1). Cell morphology quantification underwent the same process of initial image processing and masking of identified and filtered elements; however, images of the entire cell growth surface could be measured at once for a robust analysis. A table of the calculated parameters from regionprops() was exported for further evaluation in R Studio.

